# Prediction and validation of the three-dimensional structure of glucokinase-1 from *Phytophthora infestans*

**DOI:** 10.1101/2021.09.10.459842

**Authors:** Liara Villalobos-Piña, Ascanio Rojas, Héctor Acosta

## Abstract

According to its primary structure, the [PITG_06016] gene encodes for one of the 7 glucokinases present in *Phytophthora infestans* (*Pi*GlcK-1), the causal agent of late blight disease. Currently, there are no structural studies of any of its enzymes, being the determination of the three-dimensional (3D) structure of *Pi*GlcK-1 a necessary contribution in the deduction of its functions, its interaction with ligands, and possible regulatory mechanisms. In this work we present the first structural model obtained by *in silico* tools for *Pi*GlcK-1. For the prediction of this model, different algorithms were used to find the best annealing, refinement, and qualitative evaluation of them. A structural comparison of the predicted model with other structures of crystallized kinase enzymes allowed us to identify the regions of interaction with their classical substrates (glucose and ATP), as well as to identify the amino acid residues involved in the binding of other substrates such as fructose and ADP. In addition, we propose a possible recognition region of PPi, an activator of kinase activity that includes the GXGE motif, conserved in enzymes of the ribokinase (RK) family, which distinguishes this *Pi*GlcK-1 from a classical glucokinase. Accordingly, these findings suggest PPi-binding motif as potential targets for the development of inhibitors of *Pi*GlcK-1 activity.

## Introduction

*Phytophthora infestans* is the causal agent of late blight disease, which affects potato and tomato crops worldwide, causing significant economic losses in the production of these crops [1]. The (PITG_06016) gene codes for one of the 7 glucokinases present in this phytopathogen [2]. This glucokinase, known as *Pi*GlcK-1, is very important due to its high degree of expression precisely at the infectious stages of its life cycle [3]. *Pi*GlcK-1 was also the first *P. infestans* glucokinase to be biochemically characterized, and based on its sequence, it would belong to the glucokinase A group of the hexokinase family. Certainly, the characterization of *Pi*GlcK-1 has revealed the versatility of this enzyme in the phosphorylation of both glucose and fructose, as well as in the utilization of ATP, ADP, and PPi as phosphoryl donors [4].

These findings raise the need for further structural analyses on *Pi*GlcK-1. In this sense, a better understanding of the three-dimensional (3D) structure of this protein, as well as the spatial description of the possible binding sites to various ligands would contribute to the knowledge of potential targets for the design of inhibitors of the enzymatic activity of *Pi*GlcK-1, a key enzyme in the metabolism of *P. infestans*.

However, experimental determination of protein structures remains a costly and time-consuming challenge. In fact, to date no *P. infestans* enzyme has been structurally characterized. In contrast, bioinformatics tools offer an alternative that allows prediction of 3D protein structures by molecular dynamics and homology modeling which is faster, cheaper, and highly reliable. The reliability of these tools is based on the use of databases of solved protein structures that can serve as homology templates in the simulation and structural prediction of proteins [5].

In this work we present the first model obtained by *in silico* tools of the 3D structure of *Pi*GlcK-1 and the binding sites to its classical substrates glucose and ATP, as well as to fructose and ADP, and a possible binding site to PPi that is propose as a promising target for the design of an inhibitor of this enzyme.

## Materials and methods

### Sequence availability

The amino acid sequence of the protein *Pi*GlcK-1 was obtained from the NCBI database under accession number XP_002998228.

### Structural analysis of *Pi*GlcK-1 protein

The 3D structure of the protein *Pi*GlcK-1 was obtained by homology modeling using the Phyre2 [6], LOMETS [7] and I-TASSER [8] algorithms. Each simulation obtained was refined with Galaxy Refine [9] to the maximum allowed, in some cases up to three times. Validation of structural stability was performed by qualitative evaluation by ModFOLD6 [10] and by Ramachandran plot analysis obtained with SwissModel [11]. The topology of the *Pi*GclK-1 model was obtained with Pro-origami [12]. Visualization and editing of the models was performed in Chimera 1.15.1 [13].

### Docking

Molecular docking was carried out in SwissDock [14] using the refined *Pi*GlcK-1 model and substrates obtained from PubChem [15]: glucose (107526), fructose (2723872), ATP/Mg^+2^ (5957), ADP/Mg^+2^ (6022), and PPi/Mg^+2^ (644102). In each case, the best energetic fits, such as model interaction energy of the complex bound to *Pi*GlcK-1 and *fullfitness* values, were evaluated. Model visualization and editing was performed with Chimera 1.15.1 [13].

## Results

### Tertiary structure modeling

The models obtained from the structure of *Pi*GlcK-1 in three different algorithms (Phyre2, LOMETS and I-TASSER) were highly satisfactory, obtaining overall scores greater than 0.4 and a p-value cut-off of less than 0.001 which are considered very good (Table 1).

**Table 1.** Models built by various servers and their evaluation results.

| *Model* | *Favored region (%)* | *Confidence and p-value* | *Global model quality score* |
| --- | --- | --- | --- |
| Phyre2 |  |  |  |
| Before refinement | 91.50 | 4.42 E-7 | 0.6220 |
| After refinement<sup>a</sup> | 97.95 | 1.93 E-7 | 0.6412 |
| LOMETS |  |  |  |
| Before refinement | 93.45 | 5.14 E-8 | 0.6718 |
| After refinement<sup>b</sup> | 96.30 | 5.22 E-8 | 0.614 |
| I-TASSER |  |  |  |
| Before refinement | 81.60 | 3.56 E-8 | 0.6803 |
| After refinement<sup>c</sup> | 96.01 | 2.83 E-8 | 0.6854 |

<sup>a</sup> 2 time, <sup>b</sup> 3 time, <sup>c</sup> 3 time

In each case, the models were refined until the best structure was obtained, according to the Ramachandran graph, with the model yielded by Phyre2 being the most suitable due to its higher stereochemical quality, with 97.95% of the amino acids in Ramachandran regions favored, making it the potentially reliable and good quality 3D model of the *Pi*GlcK-1 protein for this work (Figure 1).

**Fig. 1.**
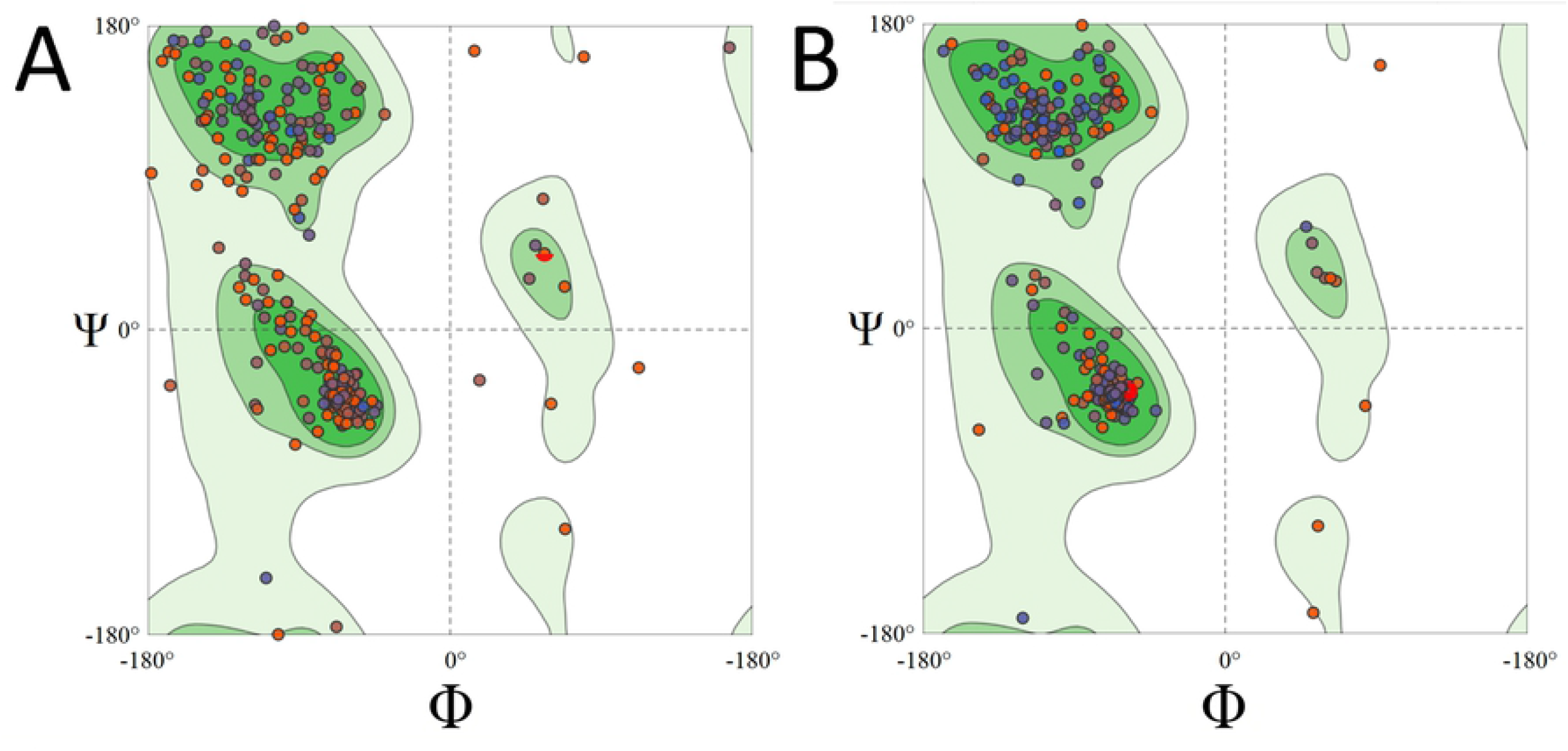
Ramachandran plot of *Pi*GlcK-1 model for Phyre2 before (A) and after (B) refinement.

The predicted model indicated that the monomeric structure was composed of two domains, a small one including residues 7 to 128, and a large domain formed by residues 135 to 343. Both domains were consecutively labeled from N-terminal to C-terminal and linked by a hinge (Figure 2).

**Fig. 2.**
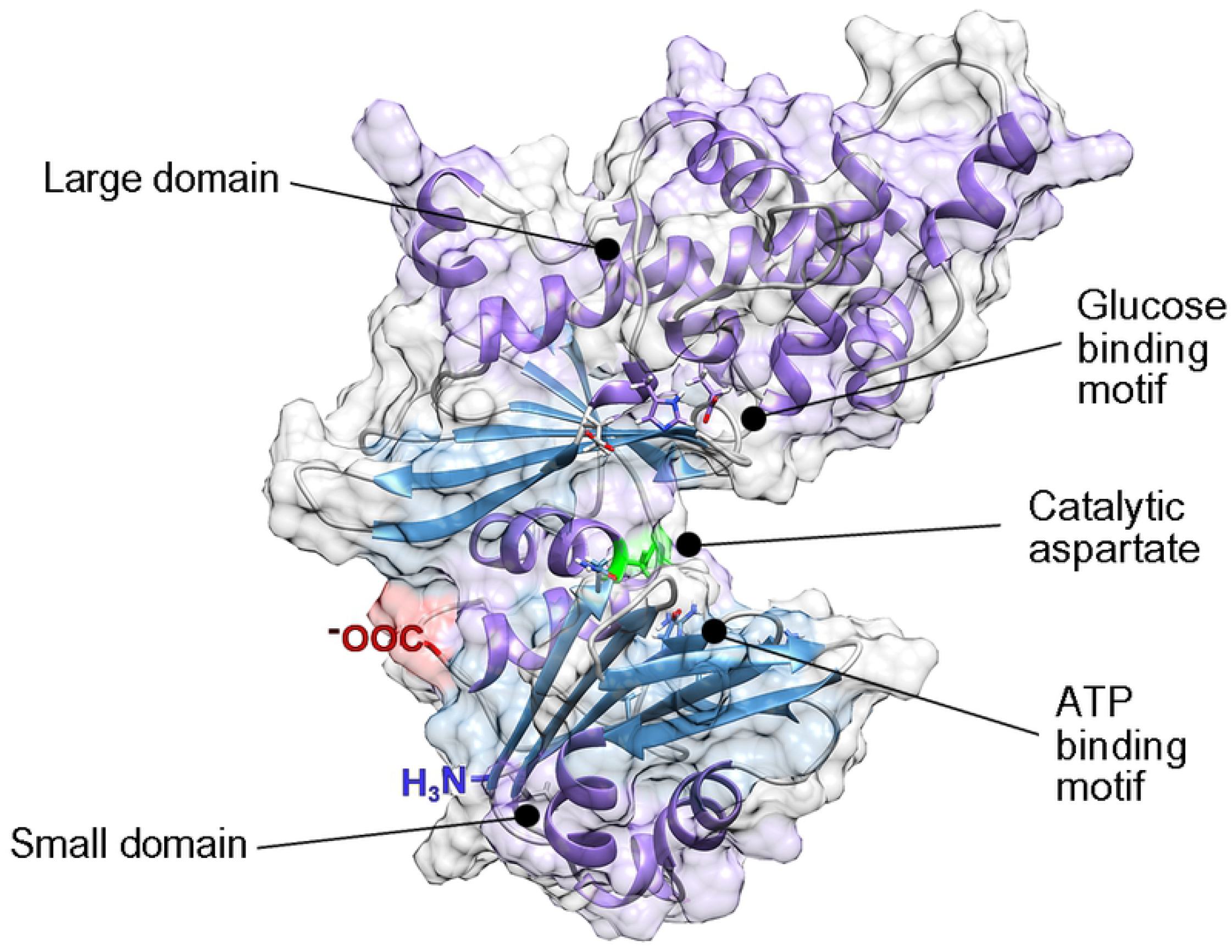
*Pi*GlcK-1 model 3D predicted by Phyre2. *Pi*GlcK-1 monomer showing the glucose binding motif in the major lobe, the ATP binding motif in the minor lobe and the catalytic aspartate binding motif in the hinge region.

The topology obtained by Pro-Origomi (S1Figure) showed that the small domain consists of 4 α-helices and 7 β–sheet distributed in two mixed central sheets; one of them of 5 stranded β (β1, β2, β3, β4 and β7) with β2 anti-parallel to the rest and another one formed by 2 β–sheet (β5 and β6) anti-parallel to each other. These sheets are flanked by 4 α–helix (α1, α2, α3 y α4).

On the other hand, the large domain consists of 10 α-helices and 6 β–sheet organized in a single mixed central sheet containing all 6 chains β (β8, β9, β10, β11, β12 and β13) with β8 and β10 anti-parallel to the rest. This sheet is flanked by 10 α–helices (α5, α6, α7, α8, α9, α10, α11, α12, α13 and α14) (S1).

### Molecular docking studies

In order to know the active site of the *Pi*GlcK-1 a docking was performed with its classic substrates glucose and ATP-Mg^2+^. This analysis showed that the glucose binding site is in the large domain while the ATP binding site is in the small domain; both molecules interact with an extensive network of hydrogen bonds within the active site (Figure 3). These results were compared with those obtained by crystallography for GlcK from *Escherichia coli* (EcGlcK) (PDB: 1SZ2), GlcK from *Trypanosoma cruzi* (TcGlcK) (PDB: 2Q2R), and the 3D model of the complex GlcK-Mg^2+^-ATP-glucose (GMAG) of human [16].

**Fig. 3.**
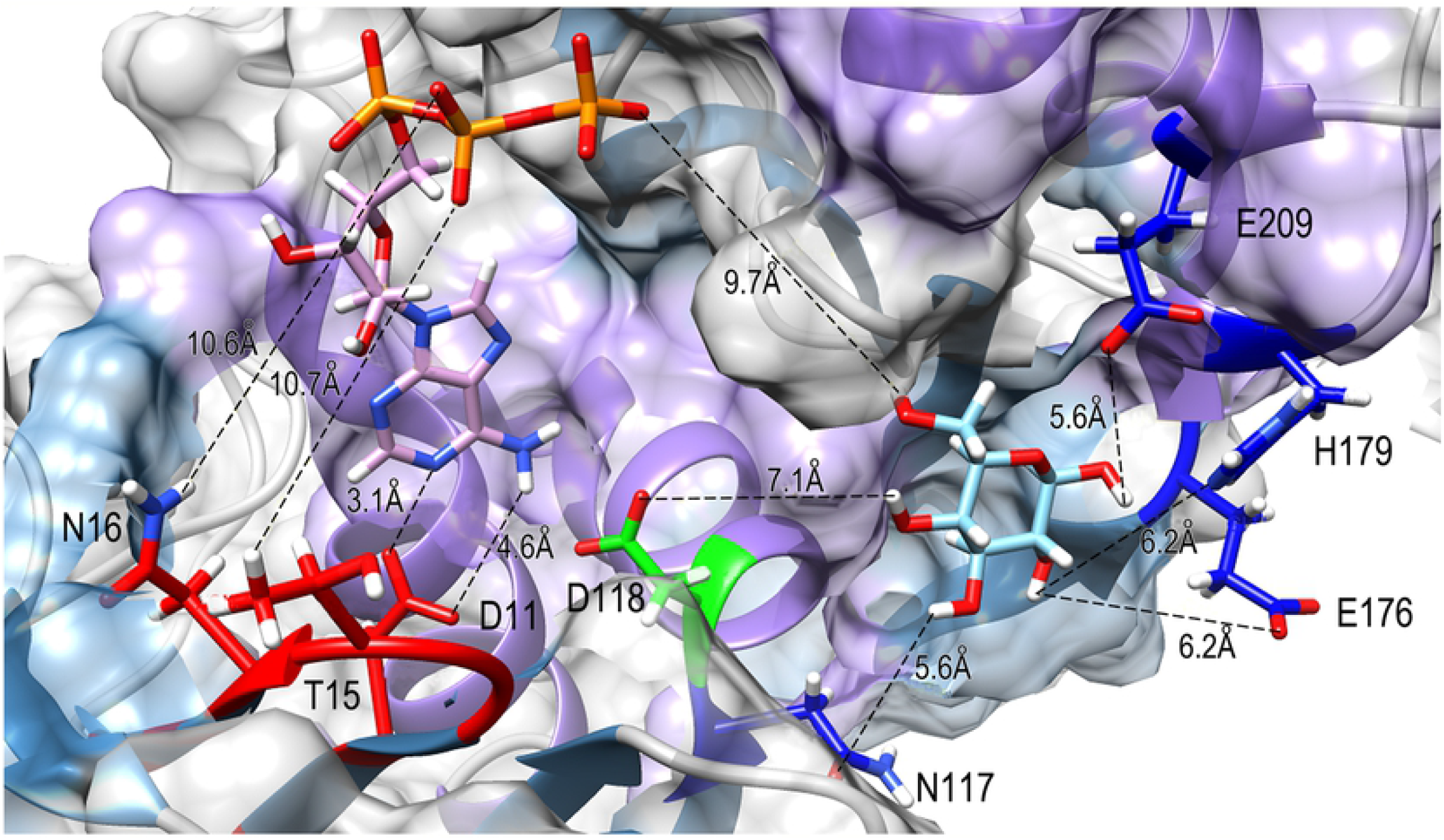
The binding site region between *Pi*GlcK-1 and glucose and ATP. Docking result using Swiss Dock. Amino acids in *Pi*GlcK-1 involved in the interaction with their ligands are highlighted in red (ATP) and blue (glucose). Catalytic aspartate is highlighted in green.

In this sense, the hydrogen bonds with the best values of ΔG y *fullfitness* were obtained from the residues Glu 209; Glu 176, His 179; Asn 117, and Asp 118 with atoms O1; O2; 03 and O4 of glucose with a ΔG of −5.27 and a *fullfitness* value of −1746.84 (Figure 3). Whereas for the β-phosphoryl group of ATP, the best values were obtained with the amino acids Thr 15 and Asn 16, and the γ-phosphoryl interacting with the O6 atom of glucose with an energy of −11.69 kcal/mol and a *fullfitness* value of −2187.34 (Figure 3). Notably, all the interacting residues described above are highly conserved in group A GlcKs from diverse organisms [4].

### Fructose-binding site

We have previously demonstrated experimentally that *Pi*GlcK-1 is able to phosphorylate other sugars such as fructose [4]. To identify a possible binding site for this hexose, we compared the fructose binding site with that of glucose, obtaining that this sugar interacts with the same amino acid residues described for glucose in this work. Fructose binding, as for glucose, occurred in the large domain and hinge region of *Pi*GlcK-1 with an energy of −5.742 kcal/mol and a *fullfitness* value of −1728.80 (Figure 4A).

**Fig. 4.**
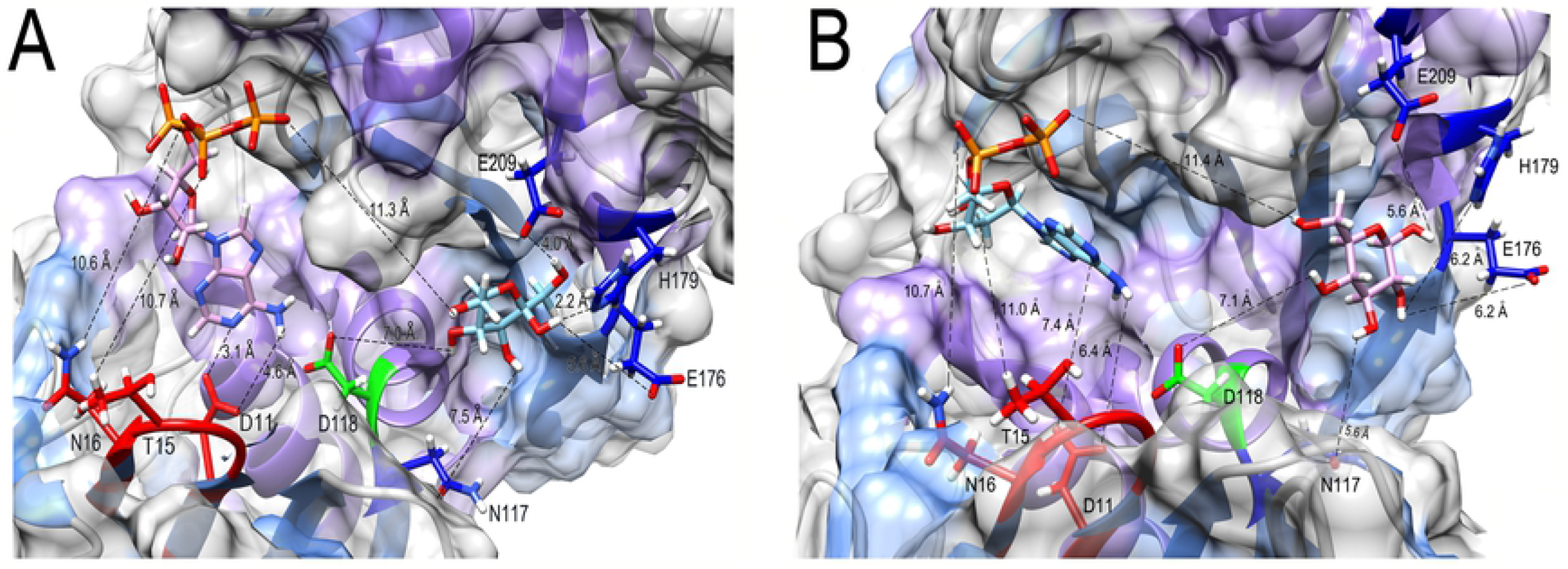
Docking result using Swiss Dock for fructose (A) and ADP (B) to *Pi*GlcK-1. (A) Amino acids in *Pi*GlcK-1 involved in the interaction with their ligands are highlighted in red (ATP) and blue (fructose). (B) Amino acids in *Pi*GlcK-1 involved in the interaction with their ligands are highlighted in red (ADP) and blue (glucose).

### Identification of ADP and PPi-binding site

One of the particularities of *Pi*GlcK-1 is to use ADP or PPi as a phosphoryl donor for the formation of hexose-6-phosphate. Additionally, PPi proved to be an efficient activator of *Pi*GlcK-1 when the phosphoryl donors are ATP or ADP [4]. Initially, we determined the binding site of ADP, by performing a docking of this ligand to *Pi*GlcK-1, finding that it indeed interacts with the same residues present in the small domain described for ATP with an energy of −10.59 kcal/mol and a *fullfitness* of −2041.78 (Figure 4B).

For its part, the best docking obtained for *Pi*GlcK-1-PPi was located in a region other than the ATP-binding and ADP-binding region, this being the larger domain with an energy of −5.425 kcal/mol and a *fullfitness* of −2150.14 (Figure 5A). Interestingly, the amino acids involved in this interaction, Lys 282, Thr 180 and Arg 210 (Figure 5B), have already been reported for the PPi-dependent kinase from *Thermotoga maritima* [17]. Markedly, *Pi*GlcK-1 also possesses the conserved GXGD(E) motif consisting of Gly 156, Leu 157, Gly 158 and Glu 159. This motif has been described in the ADP-dependent kinases of the Ribokinase family (RK) [18,19] in a region very close to the previously described PPi-binding motif, thus suggesting that residues involved in phosphoryl binding in RKs could also be participating in interaction with PPi in *Pi*GlcK-1. However, although this motif is described to bind ADP, in our case this is not possible since the ADP would be in a region other than the hinge region.

**Fig. 5.**
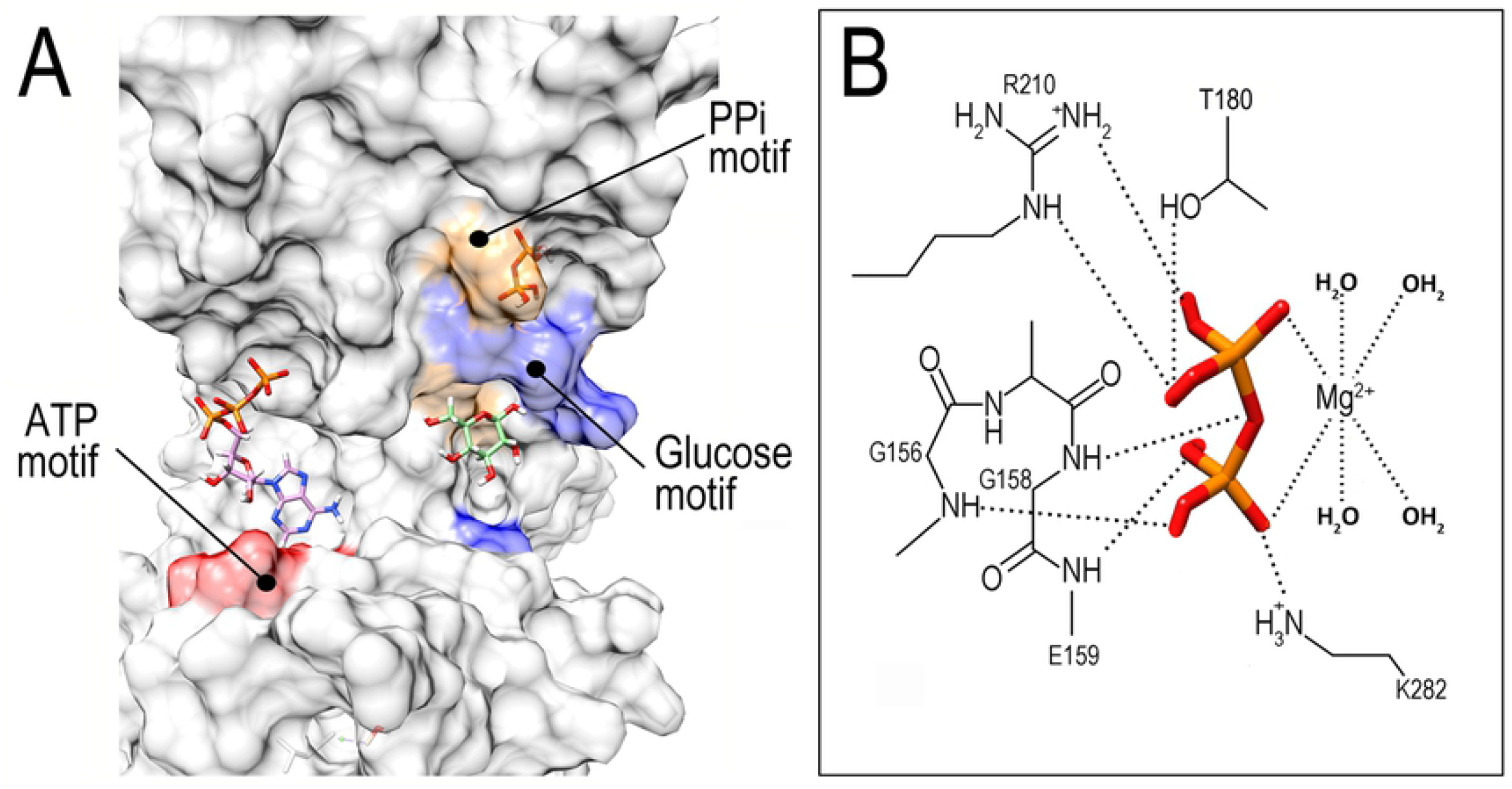
Docking result using Swiss Dock for PPi to *Pi*GlcK-1. (A) Overview of the PPi binding site (orange) distinct from that of ATP (red) and glucose (blue). (B) Close-up of the PPi binding region showing the GXGE motif and specific amino acids involved in PPi recognition.

## Discussion

Using molecular modeling and simulation methods, we constructed the 3D structural model for *Pi*GlcK-1. This model was validated and structurally compared with kinase enzymes of the HK and GlcK A family from other organisms, verifying that *Pi*GlcK-1 folds similarly to HK from yeast [20], HK from human [16] and GlcK from *E. coli* [21], whose crystallographic structures have already been described.

With our model of *Pi*GlcK-1 we were able to verify that the monomeric structure of the enzyme is highly conserved as it is composed of a large domain and a small domain connected to each other by a hinge region; both domains are made of β-sheet, flanked by α-helices, and between both lobes a cavity where the catalytic site is located, similar to many hexose kinases from several phylogenetically unrelated organisms [16, 20, 21].

Molecular docking assays on *Pi*GlcK-1 verified that ATP and glucose do indeed interact via hydrogen bonds with the residues that form the active site of the protein, as it has been described for several hexokinases and glucokinases [20]. The glucose binding site comprises well-conserved residues in the hinge region (Asp 118, Asn 117) and the large domain (Glu 176, His 179, Glu 209). In contrast, ATP binds mainly to residues Asp 11, Thr 15, and Asn 16 of the small domain with the γ-phosphoryl group pointing toward the hinge region, as it has been reported for hexokinase and glucokinase enzymes from various organisms [20].

In each case, the calculated distances between the enzyme and its substrates were made on a rigid model, so the value of these exceeds 2 Å. However, it is worth considering the intrinsic flexibility of different GlcKs [20, 21] which has reached in *Pyrococcus furiosus* a displacement of 12 Å when comparing the protein in the presence and absence of glucose [22]. Thus, the interactions between the ligands and the amino acid residues in *Pi*GlcK-1 proposed here are highly consistent with the crystallographic structures mentioned above.

Furthermore, it has been observed that fructose from the plant, like glucose, is important for the metabolism of *P. infestans* during the establishment of infection, where its concentration in potato and tomato leaves is higher than that of glucose [23–25]. Due to the absence of a specialized fructokinase (EC 2.7.1.4) in *P. infestans* [25] this sugar must be obligatorily phosphorylated by *Pi*GlcK-1.

Therefore, having corroborated in the docking of *Pi*GlcK-1 with glucose and ATP that these were located in the active site, interacting with amino acid residues conserved for other kinases, docking was carried out for fructose as well as for the phosphoryl donors ADP and PPi.

As expected, both fructose and ADP are able to form hydrogen bonds with the same amino acid residues described for their glucose and ATP pairs, respectively, validating that these can be utilized under certain physiological conditions during the life cycle of *P. infestans*. In this case, when glucose or ATP concentrations are low, *P. infestans* could obtain energy and/or NADPH through glycolysis or the pentose phosphate pathway, which would function from the use of fructose and ADP, thus ensuring that the enzyme remains active during the establishment of infection, regardless of the fact that the availability of glucose or ATP is compromised.

On the contrary, the docking for PPi, located in the large domain, close to the hinge region but in a site different from the ATP/ADP binding site (Figure 5A), would explain its role as an activator of *Pi*GlcK-1 activity in the presence of ATP or ADP [4]. This activating role of PPi could be related to a possible stabilization of a catalytically more active conformation of the enzyme, although more assays are needed to corroborate this hypothesis.

The importance of the role played by PPi in *P. infestans* has been evidenced by several facts such as : i) *P. infestans* can produce PPi through multiple biosynthesis and hydrolysis reactions carried out in this phytopathogen [26], ii) it expresses significant levels of several enzymes relevant to its energy metabolism which are PPi-dependent, such as pyruvate phosphate dikinase (PPDK), 6-phosphofructose-1-kinase (PFK), and 6-phosphofructose-2-kinase(PFK2) [3,25,27], iii) it has been suggested that the majority of the glycolytic flux in the hyphal sporulation stage of *Phytophthora cinnamomi* occurs via PPDK [28], iv) PPDK may play a more relevant role than PYK in glycolysis of *P. infestans* due to a higher expression of PPDK relative to PYK [29], v) pyrophosphate stimulates calcium uptake in diverse organisms including *P. infestans*, which is required for its growth and development [30]

Although the true value of PPi in the metabolism of *P. infestans* remains to be clarified, it is important to consider the binding motif of this ligand when designing potential inhibitors of *Pi*GlcK-1 activity as it constitutes a promising target for attack in the search for a solution to the threat posed by late blight.

## Conclusion

In this work we have presented the first *in silico* structural model of glucokinase-1 (*Pi*GlcK-1) from the phytopathogen *P. infestans*. The resulting structure was shown to be of high quality, which allows it to be used in docking analyses for other ligands. We confirmed that the folding of this protein is similar to that of other kinases from different organisms and we were able to identify the binding sites for its substrates glucose and ATP as well as for the ligands fructose, ADP and PPi reported for the first time for a classical glucokinase of the GlcK-A group. Altogether, these findings lay the groundwork for the design of future inhibitors of *Pi*GlcK-1 enzymatic activity that would aid in the control of late blight disease and underscored the need to establish a new classification within a more diverse group of kinases, one that considers not only the primary sequence of *Pi*GlcK-1, but also its 3D structure.

## Acknowledgments

We are very grateful to experiment.com and specially to Chralie Kinsella, Brian Repez, Amy Collete and Pureum Kim for their valuable help in financing this project.

## Author Contributions

**Conceptualization:** Liara Villalobos-Piña, Ascanio Rojas, Héctor Acosta

**Formal analysis:** Liara Villalobos-Piña, Ascanio Rojas, Héctor Acosta.

**Funding acquisition:** Laboratorio de Fisiología, Universidad de Los Andes, Centro de Cálculo de la Universidad de Los Andes, Laboratorio de Enzimología de Parásitos, Universidad de Los Andes.

**Methodology:** Liara Villalobos-Piña.

**Project administration**: Ascanio Rojas.

**Supervision:** Ascanio Rojas, Héctor Acosta.

**Visualization:** Héctor Acosta.

**Writing – original draft:** Liara Villalobos-Piña

**Writing – review & editing:** Liara Villalobos-Piña, Ascanio Rojas, Héctor Acosta.

## Supplementary data

**S1.**
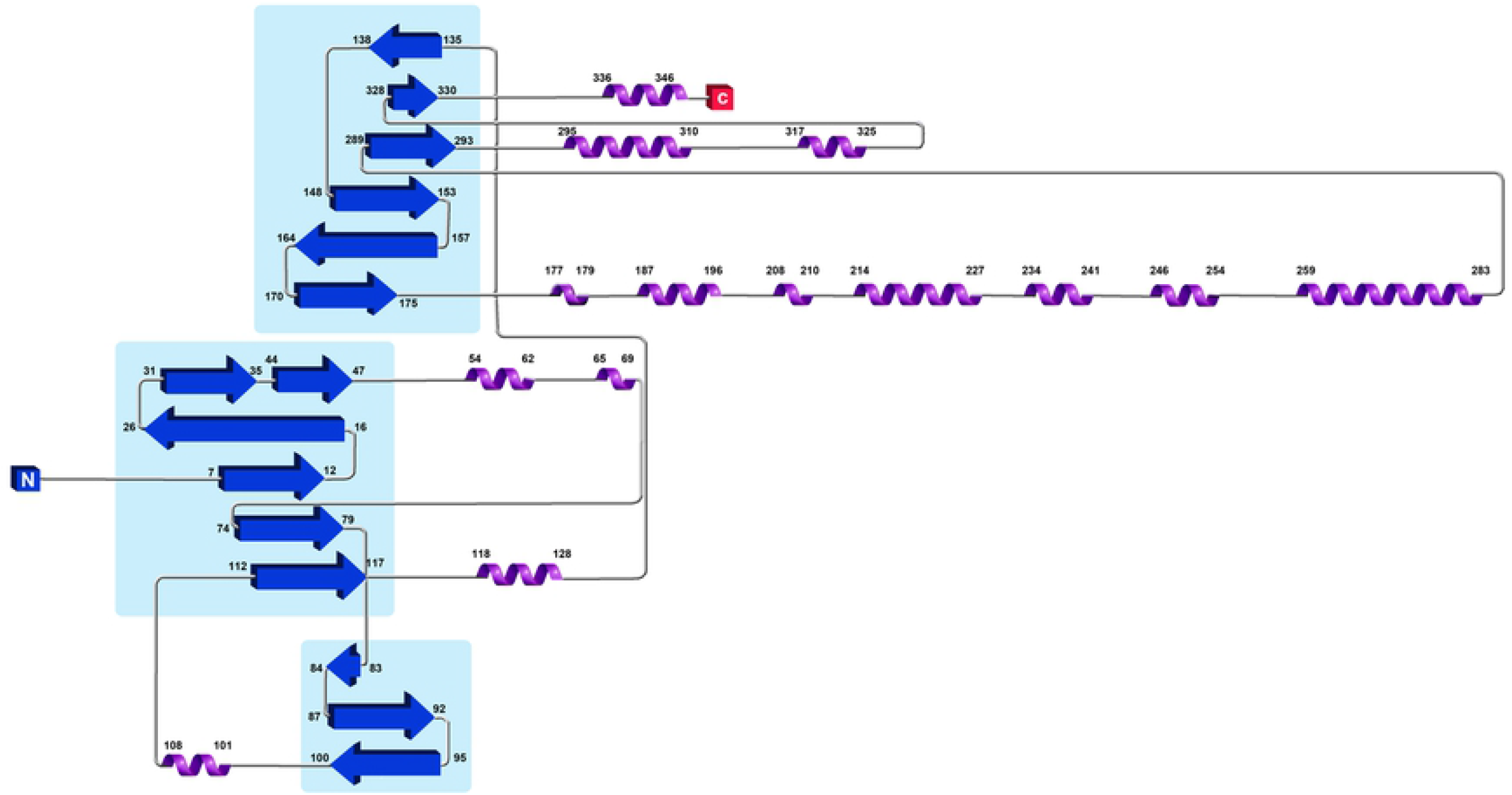
Fold topology of *Pi*GlcK-1 monomer obtained by Pro-Origami. Topology of *P. infestans* with two domains (light blue shading), composed of lamellae β-sheet (dark blue) and flanked by α-helices (purple).

